# A flexible target-specific anti-infection therapeutic platform that can be applied to different microbial species

**DOI:** 10.1101/2020.04.20.049791

**Authors:** Makoto Mitsunaga, Kimihiro Ito, Takashi Nishimura, Hironori Miyata, Yoshimitsu Mizunoe, Hisataka Kobayashi, Tadayuki Iwase

**Affiliations:** Division of Gastroenterology and Hepatology, Department of Internal Medicine, The Jikei University School of Medicine, Tokyo, Japan; Animal Research Center, School of Medicine, University of Occupational and Environmental Health, Kitakyushu, Japan; Department of Bacteriology, The Jikei University School of Medicine, Tokyo, Japan; Molecular Imaging Program, Center for Cancer Research, National Cancer Institute, NIH, Bethesda, USA; Research Center for Medical Sciences, The Jikei University School of Medicine, Tokyo, Japan

## Abstract

The emergence of new microbial pathogens, including drug-resistant strains, complicates treatment, thereby threatening global health. We demonstrated a photoimmuno-antimicrobial strategy (PIAS) that eliminated antibody-targets using a photo-activated anti-pathogen antibody generating mechanical stress that damaged the target’s binding sites. PIAS is effective against many pathogens, including methicillin-resistant *Staphylococcus aureus* (MRSA), the fungal pathogen *Candida albicans*, and viral particles irrespective of their species or drug-resistance status. Animal experiments demonstrated that PIAS saved mice from fatal infections; microbiome and histochemical analyses indicated no apparent effect on normal host microflora and tissues in PIAS-treated mice. Resistance to PIAS was not observed during the eight years of this study. As a new type of anti-infection therapy, PIAS may contribute to advances in anti-infection strategies.

## Introduction

Discovery and development of novel antimicrobial agents and strategies have a significant impact on medicine, protecting numerous lives from fatal infections (ref. 1). However, the emergence of drug-resistant pathogens has complicated the use of antimicrobial agents (refs. 2–4). Further, emerging infectious diseases such as COVID-19 infection, without available antimicrobials and strategies, are a great threat to global health.

At present, anti-infective platforms that can be applied to various pathogens irrespective of their species or drug-resistance status and that can specifically eliminate the targeted pathogens without affecting the host microflora or tissues are strongly needed in clinical settings. However, the development of new antimicrobials has reached a saturation point (refs. 1–3). To overcome this situation, different perspectives such as those generated from even seemingly irrelevant interdisciplinary fields are necessary.

The phthalocyanine derivative photoplastic probe IRDye700DX (IR700) can change its structure by near-infrared (NIR) illumination and generate mechanical stress through structural change (ref. 5). We previously evaluated the anticancer properties of this probe using an antibody against human epithelial growth factor receptor type 2 (refs. 6). Recently, it was reported that this anticancer effect depended on mechanical cell damage to the binding sites via NIR-induced structural change in a different manner than photodynamic therapy or conventional antimicrobials, including antibiotics (ref. 5). The efficacy and safety of anticancer therapy using this probe were demonstrated in clinical trials for patients with recurrent head and neck cancer (ref. 7).

During our cancer research, we conceived the potential of the antimicrobial effect of this probe. However, the verification of this concept was difficult as almost all microbial pathogens have very thick cell walls and because the work is from rather different fields. To achieve our aim of potentially changing the current threat of emerging infectious disease and drug-resistant pathogens to global health, we organized a team of researchers from multidisciplinary fields and addressed this issue from various aspects, including microbiology and immunology, infectious disease, and drug discovery.

In the present study, we designed a new, flexible anti-infective therapeutic platform that can specifically act on antibody-targets irrespective of their species or drug-resistance status, by using the IR700 photoplastic probe conjugated to an antibody against target pathogens. This antibody can disrupt the target’s binding sites through the transmission of the mechanical stress generated by NIR-induced structural changes in the probe to the pathogen’s epitopes (Supplementary Figure 1). To verify the concept, we designed a photoimmuno-antimicrobial strategy (PIAS) based platform and evaluated whether PIAS specifically eliminated antibody targets, including methicillin-resistant *Staphylococcus aureus* (MRSA), the fungal pathogen *Candida albicans* and viral particles. Further, we examined whether PIAS eliminated the target pathogen without effecting the normal mouse microflora and tissue using microbiome and histochemical analyses. In addition, we examined whether resistance to PIAS emerged during the study period (2013-present).

## Results

### Effect of PIAS on the bacterial pathogen *S. aureus*

First, we focused on *S. aureus* (ref. 8), a causative agent for skin infections such as folliculitis and impetigo to severe infections such as femoral head necrosis and fatal sepsis. In addition, methicillin-resistant *S. aureus* (MRSA) (ref. 9), which displays multi-drug resistance, is a cause of hospital-acquired infection that makes infection therapy difficult. Further, no vaccines against *S. aureus* have been developed despite considerable studies for several decades. *S. aureus* colonises the nasal cavity of approximately 30% of humans (ref. 8). Immunity against *S. aureus* is acquired; nonetheless, no eradication of the pathogen has been observed and the underlying mechanisms have not been determined (ref. 10). Given that anti-*S. aureus* antibodies bind the pathogen (although they exhibit no effects on *S. aureus* growth), we assumed that PIAS using antibodies can exhibit an antimicrobial effect on the pathogen because the ability to bind, but not inhibit, is critical for PIAS.

To begin, we generated an *S. aureus*-targeted conjugate using an anti-*S. aureus*-cell-wall-epitope monoclonal antibody (SA mAb), which exhibits no apparent effect on *S. aureus* growth and IR700 probe (SA-IR700 conjugate, SA-IR700). Then, we confirmed the binding of the probe to SA mAb by detection of the mAb and the fluorescence of the probe by sodium dodecyl sulphate polyacrylamide gel electrophoresis (SDS-PAGE) (Supplementary Figure 2). We then evaluated the ability of SA–IR700 to bind to target *S. aureus* cells through flow cytometry, fluorescence microscopy, and electron microscopy. In the flow cytometric analysis, a strong IR700 signal was observed in all the *S. aureus* strains tested (shown by red lines; Supplementary Figure 3A), whereas no signal was observed in the non-target *Escherichia coli*. When *S. aureus* cells were pretreated with SA mAb prior to treatment with SA-IR700, the fluorescence signals of the conjugate were decreased (shown by black lines; Supplementary Figure 3A). In addition, fluorescence microscopy detected fluorescence signals in all tested *S. aureus* strains but not in *E. coli* (Supplementary Figure 4B). Further, immune-gold staining and field emission scanning electron microscopy (FE-SEM) analysis demonstrated that the conjugates bound to the targets, as shown by the colloidal gold secondary antibody (top panel; Figure 1A). These data indicate that the conjugates can bind to the target based on the immune-specificity of the mAb used.

**Figure 1.**
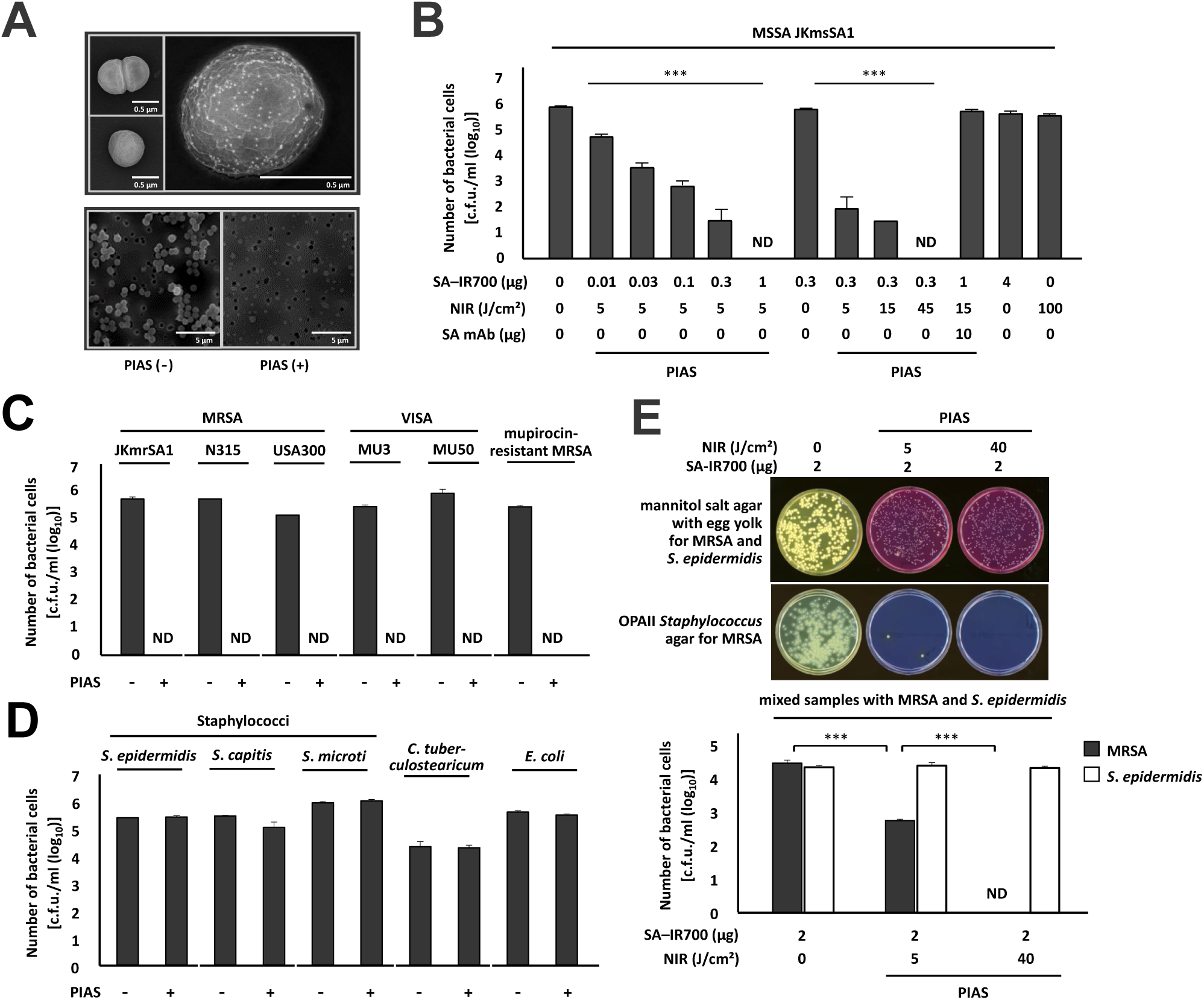
Bactericidal effect of photoimmuno antimicrobial strategy (PIAS). **A**. FE-SEM analysis of SA–IR700 conjugate-bound *S. aureus* cells subjected to immunogold staining (top images). Non-treated (left top in top images) and colloidal gold IgG-treated cells (left bottom in top images). (right in top images) Bright dots on bacterial cell treated with SA–IR700 and colloidal gold secondary antibody indicate the presence of conjugates. SEM analysis for PIAS-damaged *S. aureus* cells (bottom images). Non-PIAS-treated (left top in bottom images) and PIAS-treated cells (right in bottom images). **B.** Bactericidal effect of PIAS on *S. aureus* cells. **C** and **B.** Bactericidal effect of PIAS on bacterial cells of various drug-resistant *S. aureus* strains (**C**) and on those of non-target bacteria (**D**). **E.** Target-selective bactericidal effect of PIAS on mixed samples with MRSA and the non-target *S. epidermidis*. Yellow or pink colonies in the panel **E** are *S. aureus*/MRSA or *S. epidermidis* colonies, respectively. Representative images are shown. c.f.u., colony-forming units; ND, not detected. Data are representative of at least three independent experiments, and mean values are shown. Error bars represent standard deviation from triplicate samples. (**B**) One-way analysis of variant (ANOVA) and (**E**) ANOVA with Sidak test. \*\*\**P* < 0.0001.

To investigate the bactericidal effect of the conjugate, PIAS was used along with SA–IR700 on *S. aureus* cells. Briefly, bacterial cells [10^6^ colony-forming units (c.f.u.)/test] harvested from the logarithmic and stationary (ref. 11) phases were treated with the conjugates (0 to 4 µg/test), followed by NIR illumination (0 to 100 J/cm^2^). The PIAS-treated bacterial cells were cultured on agar plates, and colonies were counted to evaluate the bactericidal effect.

PIAS exhibited the bactericidal activity on *S. aureus* cells in a conjugate- and NIR-dose-dependent manner and PIAS-treated pathogen cells were eradicated within several minutes of NIR illumination (5 J/cm^2^, 1 min) (Figure 1B). Upon treatment solely with SA–IR700, NIR illumination, or with NIR illumination with SA mAb instead of SA–IR700, no bactericidal effect was observed (Figure 1B). PIAS also affected bacterial cells under various conditions (Supplementary Figure 4).

PIAS performed effectively with an anti-*S. aureus* antibody against various epitopes although the antimicrobial effect varied slightly (Supplementary Figure 5), implying that the bactericidal effect of PIAS depends on the quality of the antibody and effects related to the antibody’s binding site and that the use of multiple conjugates of multiple anti-epitope antibodies enables a response to altered epitopes of the target pathogen.

Further, PIAS eliminated various drug-resistant *S. aureus* strains, including the methicillin-resistant *S. aureus* (MRSA) and vancomycin (VCM)-intermediate *S. aureus* (VISA) (ref. 12) that has very thick cell walls and displays resistance to almost all antibiotics (Figure 1C and Supplementary Figure 6). PIAS also eradicated cells harbouring a plasmid encoding the tetracycline-resistance gene (Supplementary Figure 7). These results indicated that PIAS acted on the pathogen cells irrespective of their drug-resistance status.

In non-targets, such as members of the subjects’ normal microflora, including *Staphylococcus epidermidis, Staphylococcus capitis, Staphylococcus microti, Corynebacterium tuberculostearicum*, and *E. coli*, no apparent effects of PIAS were observed (Figure 1D). PIAS selectively eradicated the target pathogen MRSA within mixed samples of MRSA and the non-target *S. epidermidis* (Figure 1E). The peptidoglycan of *S. epidermidis* is considered to be similar to that of *S. aureus*. However, lysostaphin that lyses the peptidoglycan of *S. aureus* exhibits no apparent effect on that of *S. epidermidis*, indicating that the peptidoglycan components of *S. aureus* and *S. epidermidis* vary and implying that the antibody used in this study could target such unique epitopes for the pathogen. Additionally, PIAS using non-specific trastuzumab (ref. 6) (anti-human epidermal growth factor receptor 2 antibody, Tra)–IR700 conjugate (Tra-IR700) exhibited no apparent effect on *S. aureus* mutant cells (Δ*spa*) or wild-type cells deficient in protein A, an IgG-binding cell wall-associated protein expressed by *S. aureus* (Supplementary Figure 8) (ref. 13). These results suggested that non-specific binding effect did not substantially contribute to the bactericidal effect, and that structural proteins such as peptidoglycan epitopes may be more effective targets for PIAS compared to non-essential cell-surface proteins.

PIAS acted on bacterial cells in the stationary phase and the growing phases, whereas β-lactam drugs such as penicillin act only on growing cells. After 30 passages, PIAS acted on new bacterial strains in the same manner as on the parental strain (Supplementary Figure 9), and no strains became resistant to PIAS during the study period. PIAS also eradicated the pathogen cells when the experiments were performed under low-temperature conditions (4°C) and in the presence of the oxygen toxicity-detoxifying enzyme, catalase (data not shown). Notably, SEM analysis of PIAS-induced damage to pathogens showed no intact cells (bottom images; Figure 1A). Based on minimal cell debris found in PIAS-treated samples, we speculate that the peptidoglycan-lysing enzymes of the pathogen cells may be involved. However, our data suggested that PIAS can act on pathogens differently than antimicrobials (ref. 14) or photodynamic therapy (ref. 15). These data indicate that PIAS specifically and promptly damages target cells when conjugate binding is followed by NIR illumination, suggesting that compared to antibiotics, induction of resistance against PIAS is unlikely.

Next, the *in vivo* effect of PIAS was investigated using a rat model of MRSA colonization (ref. 16), and mouse models of MRSA-intraperitoneal (ref. 17) and -thigh infections (ref. 18) (Figure 2).

**Figure 2.**
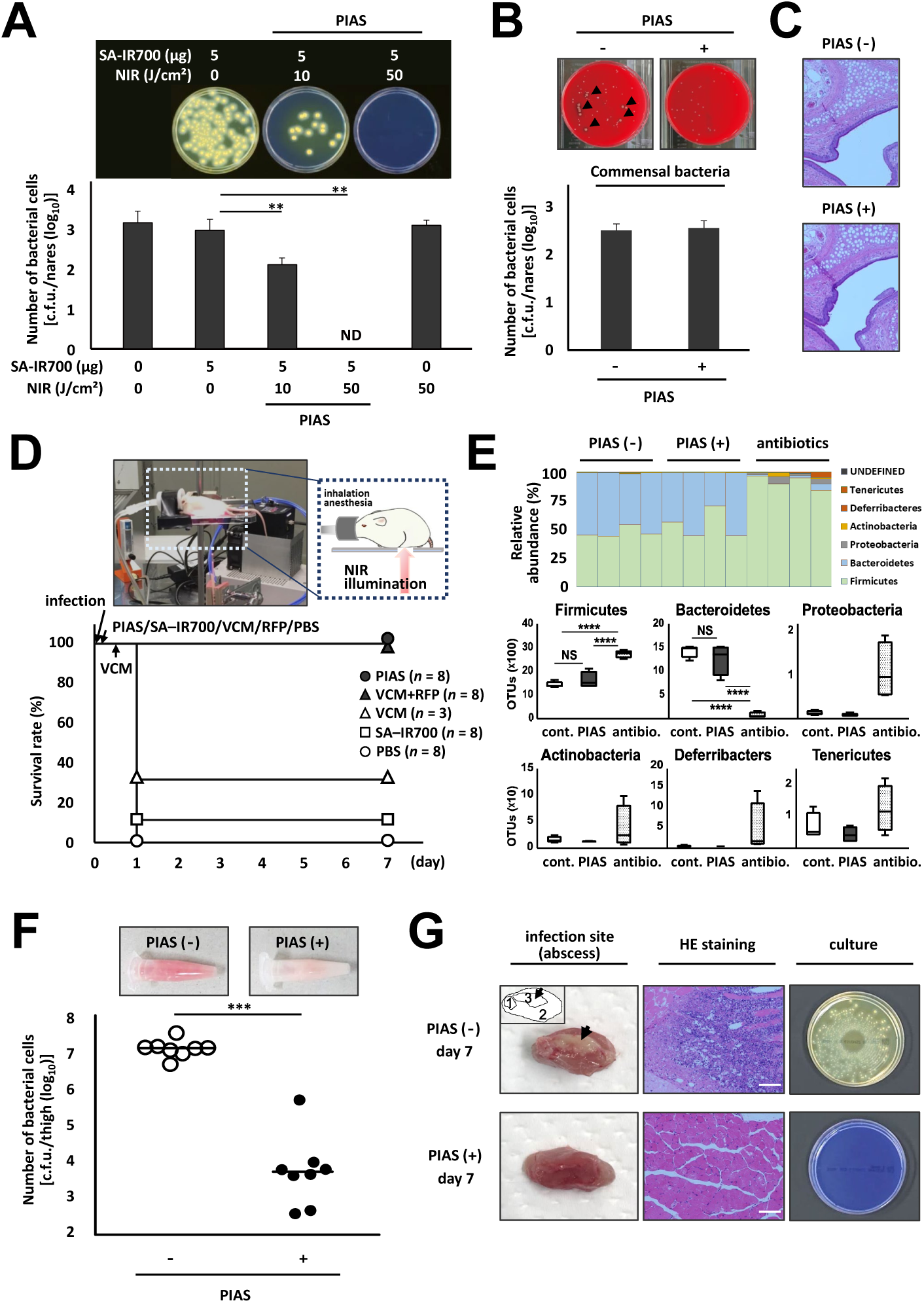
*In vivo* effect of PIAS. **A**–**C**. Cotton rats (*n* = 6 each group) colonized by MRSA JKmrSA1 were treated with PIAS. PIAS conditions: SA–IR700, 0–5 µg/rat; NIR illumination, 0–10 J/cm^2^. The samples were cultured on the MRSA-selection medium OPAII (**A**) and the non-selection medium TSA (**B**), and colonies were counted to evaluate the bactericides effect of PIAS *in vivo*. Data are summarized from two independent experiments (**A**). Arrow heads in culture images in **B** indicate *S. aureus* colonies; *S. aureus* colonies produce hemolytic plaques on TSA. **C.** Histological analysis was conducted on the nasal tissue of rats treated with or without PIAS. Representative images are shown. **D** and **E.** Effects of treatments of PIAS and conventional antibiotics on a mouse model of intraperitoneal infection (**D**) and host’s normal intestinal microflora (**E**), respectively. Mice intraperitoneally administered MRSA cells (2 × 10^8^ c.f.u./mouse), a VCM-sensitive strain, were treated with PIAS, SA–IR700, antibiotics (VCM or RFP + VCM) or PBS. PIAS conditions: SA–IR700, 50 µg/mouse; NIR illumination, 100 J/cm^2^. Antibiotics treatments: (i) VCM at 0.1 mg/mouse twice a day, (ii) RFP at 2 mg/mouse once a day and VCM at 0.5 mg/mouse twice a day. Control groups: SA–IR700, 50 µg/mouse; PBS, 0.1 ml. **D.** Effects of PIAS and antibiotics treatment on survival of MRSA-infected mice. Data are merged from two different experiments (PIAS and antibiotics treatments). The image and cartoon depict a mouse under treatment with aPIT. **E.** 16S rRNA-targeting metagenome analysis on bacterial phyla. cont, control (PBS); antibio, antibiotics (VCM + RFP). Box elements: center lines, median; box limits, upper and lower quartiles; whiskers, minimum and maximum values. **F** and **G.** Effect of PIAS on a mouse model of MRSA-thigh infection. MRSA cells (10^7^ c.f.u./thigh) injected into the right thigh muscle of mouse (*n* = 8 each group) were treated with PIAS. PIAS conditions: SA–IR700, 50 μg/mouse; NIR illumination, 50 J/cm^2^. **F.** The homogenized thigh samples (day 1; top images) were cultured on OPAII, and colonies were counted. Horizontal bars indicate median. **G.** On the thigh samples at day 7, visual (right images) and histochemical (middle) analyses and bacterial culture (right images) were performed; arrows and the cartoon (top left) depict the infection site of the thigh (1, femur; 2, thigh muscle; 3, abscess). Scale bars indicate 100 μm. HE, hematoxylin and eosin. Representative images are shown (**A**–**C, F**, and **G**). c.f.u., colony-forming units; ND, not detected. Mean values are shown. Error bars represent standard deviation from triplicate samples. one-way analysis of variance (ANOVA) with Dunnett’s test (**A**), two-tailed unpaired Student’s *t*-test (**B** and **F**) and one-way ANOVA with two-stage step-up method of Benjamini, Krieger and Yekutieli (**E**). \*\**P* < 0.01, \*\*\**P* < 0.001, \*\*\*\**P* < 0.0001.

Cotton rats, which are used in investigating *S. aureus* colonization (ref. 16), were used to evaluate the elimination of MRSA nasal colonisation. Rats were first confirmed to be non-*S. aureus* carriers and capable of retaining MRSA colonization during the study period (at least 2 weeks) after intranasal instillation with 1 × 10^6^ c.f.u. of MRSA JKmrSA1 cells. Animals passing this assessment were treated with PIAS. Seven days after intranasal instillation of the pathogen cells, cotton rats were intranasally administered SA–IR700 using a pipette (5 μL, 5 μg/rat), followed by NIR illumination. Subsequently, nasal samples were harvested and cultured on OPAII *staphylococcus* agar (OPAII) and trypticase soy agar with sheep blood (TSA); OPAII is a selection medium for MRSA (the targeted pathogen), and TSA is a non-selective medium.

Consistent with *in vitro* studies, PIAS eradicated MRSA from the rat nasal tract (Figure 2A); the rat commensal bacteria were not eradicated (Figure 2B and Supplementary Figure 10). Careful monitoring of the PIAS-treated animals revealed no physical changes, and no significant damage was found in the histological analysis of rat nasal tissue (Figure 2C). Additionally, no apparent effect of PIAS was observed on the non-target fibroblast cells (Supplementary Figure 11).

The effect of PIAS, in principle, partly depends on the amount of light available to the targets. Therefore, to determine its effectiveness against internal pathogens, a mouse model of intraperitoneal infection (ref. 17) with bacterial cells (10^8^ c.f.u./mouse) of MRSA JKmrSA1 was used. In this experiment, SA–IR700 (administered intraperitoneally) was used in both the test and control groups considering the opsonisation of internal target pathogens by the anti-pathogen antibody.

Mice treated with PIAS (external NIR illumination, without shaving their fur) survived (Figure 2D), whereas mice treated with PBS died (p > 0.0001) (Figure 2D). Upon treatment solely with SA–IR700 (without NIR illumination), one mouse survived, suggestive of the opsonisation effect (Figure 2D). To investigate the effects of PIAS treatments and conventional antibiotics on the host’s normal intestinal microflora, additional experiments with antibiotic treatment [a twice-a-day regimen of VCM or VCM and rifampicin (RFP, once a day)] were performed (Figure 2D); VCM and RFP were administered intraperitoneally and orally, respectively. Using faecal samples of PIAS-treated or antibiotic (VCM and RFP)-treated mice, 16S rRNA-targeting metagenome analysis was performed (refs. 19, 20). The analysis indicated that the antibiotic treatment affected the microflora whereas PIAS did not (Figure 2E and Supplementary Figure 12), suggesting that PIAS acted selectively on the target pathogen without an apparent effect on non-targets such as the host’s normal microflora.

Further, the effect of PIAS on a mouse model of MRSA-thigh infection (ref. 18) was investigated to evaluate the bactericidal effect of PIAS on pathogen cells in deep focus in a tissue. Briefly, mice injected with MRSA JKmrSA1 (10^7^ c.f.u./thigh) were treated with PIAS (day 0); SA–IR700 was intramuscularly administered, and external NIR illumination was administered to the thigh without shaving their fur. Thighs were harvested on day 1, homogenized, and cultured on OPAII. Additionally, histological analysis and bacterial culture were performed on day-7 thigh samples.

PIAS exhibited a bactericidal effect on MRSA cells in the thigh (Figure 2F). An apparent hyperaemia was observed in the homogenized thigh samples of non-PIAS treated mice but not in those of PIAS-treated mice (top images; Figure 2F). In day-7 samples of non-PIAS treated mice (Figure 2G), apparent abscess (left images) and inflammatory cell infiltration (middle images) were observed; MRSA cells were persistently detected in non-PIAS treated mice but not in PIAS-treated mice (right images).

Various drug-resistant *S. aureus* strains against anti-MRSA drugs such as linezolid and daptomycin in addition to VCM, which are the few drugs clinically available for MRSA infections (refs. 8), have emerged, thereby complicating anti-infective therapy (refs. 1–3). In addition, the use of antimicrobial agents can lead to dysbiosis by disrupting the normal host microflora (ref. 21) and damaging host tissues (ref. 22). The present study demonstrates the target-specific effect of PIAS, irrespective of their drug-resistant status.

### Effect of PIAS on the fungal pathogen *C. albicans*

We assumed that PIAS could be applied to a broad range of microbial pathogens (such as bacteria, fungi, and viruses) based on the specificity of the antibody used. To verify the potential of PIAS in eliminating fungal pathogens, we investigated whether PIAS works in *Candida albicans*, a pathogenic fungus that causes various diseases from superficial to severe systemic infections (ref. 23, 24). Then, we used an anti-*C. albicans* (CA) mAb to generate CA–IR700 conjugates (CA–IR700).

Consistent with the bactericidal effect of PIAS on *S. aureus*, PIAS using CA–IR700 was found to eradicate *C. albicans* (Figure 3A). Further, PIAS eliminated drug-resistant *C. albicans* strains such as azole-drug (fluconazole, itraconazole, and voriconazole; these are commonly used as anti-fungal agents) and flucytosine-resistant strains (Figure 3B), similar to non-drug-resistant *C. albicans*. In contrast, PIAS did not work on the non-targeted, non-pathogenic fungi such as *Candida stellate* (Figure 3C).

**Figure 3.**
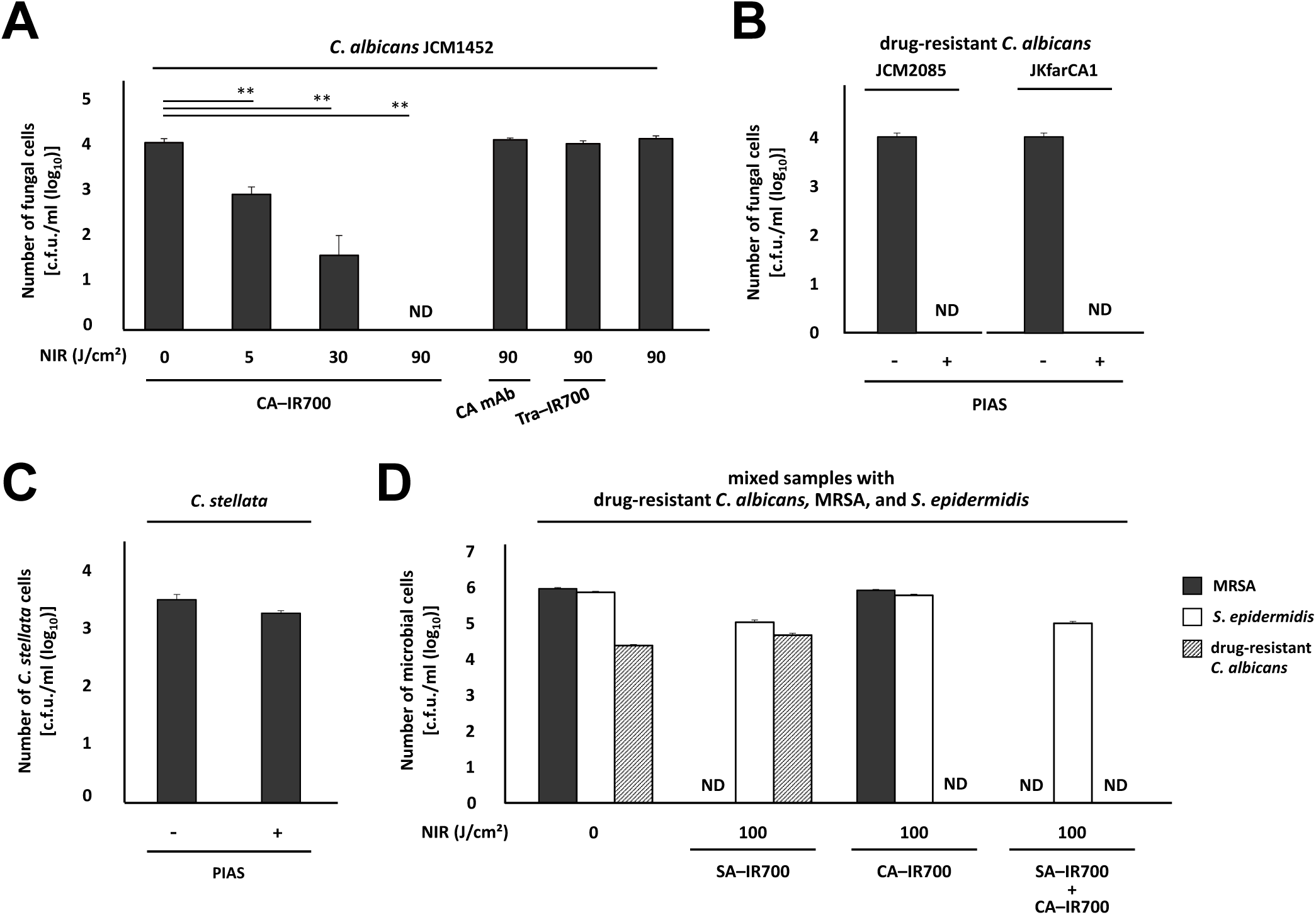
Fungicidal effect of PIAS. **A**–**C.** Effects of aPIT using CA–IR700 on *C. albicans* (**A**), drug-resistant *C. albicans* (**B**), and the non-target, non-pathogenic fungus *C. stellata* (**C**). **D.** Target-selective microbicidal effect of PIAS using CA–IR700 and/or SA–IR700 on a mixed sample with drug-resistant *C. albicans*, MRSA, and *S. epidermidis*. PIAS conditions: CA–IR700 (20 µg/test) and/or SA–IR700 (1 µg/test); NIR illumination, 100 J/cm^2^. c.f.u., colony forming units; ND, not detected. Data are representative of at least three independent experiments, and mean values are shown. Error bars represent standard deviation from triplicate samples. One-way analysis of variant with Dunnett’s test. \*\**P* < 0.01.

We also investigated the target specificity of the eradication effect of PIAS against dual conjugates using a mixed sample with drug-resistant *C. albicans*, MRSA, and *S. epidermidis*. PIAS using SA–IR700 or CA–IR700 eradicated MRSA or *C. albicans* in the mixed sample, respectively (Figure 3D). PIAS using both SA–IR700 and CA–IR700 eradicated both MRSA and *C. albicans* in the mixed sample (Figure 3D). The non-target *S. epidermidis* was not eradicated in any of the cases (Figure 3D).

These experiments indicated that using multiple conjugates against multiple pathogens, PIAS selectively eradicates the targeted pathogen(s), regardless of their species or drug-resistant status, even in the presence of non-targeted microbes.

### Effect of PIAS on viral particles and viral infection

Finally, we investigated the potential of PIAS against viral infections. We used bacteriophage T7, a bacteria-lysing virus, for this experiment owing to its high reproducibility (ref. 25). For PIAS on the phage, we used a mAb against the capsid epitope of T7 to generate T7–IR700 conjugates (T7–IR700). The phage was subjected to PIAS using the conjugates. As a control, T7 phage was treated with solely T7– IR700 (without NIR illumination) to equalize the interference of the conjugates on the infection of the phage to *E. coli* cells. These treated phage samples were mixed with *E. coli* cells, and bacterial colony formation was then investigated to evaluate the phage-inactivation effect of PIAS.

In the PIAS-treated sample, colonies were observed; however, no colonies were observed in the control sample (Figure 4), meaning that the PIAS-treated T7 phage was inactivated. Although the T4 phage was also subjected to PIAS using T7–IR700, no effect was observed (Figure 4), indicating that PIAS selectively affects viral targets based on the specificity of the mAb used. These results indicate that PIAS can be used as a strategy to directly inactivate a bacterial virus, and could contribute to the elimination of bacteriophages remaining in the body after phage therapies (refs. 26, 27). These results imply that PIAS can be used as a novel therapy for viral infections.

**Figure 4.**
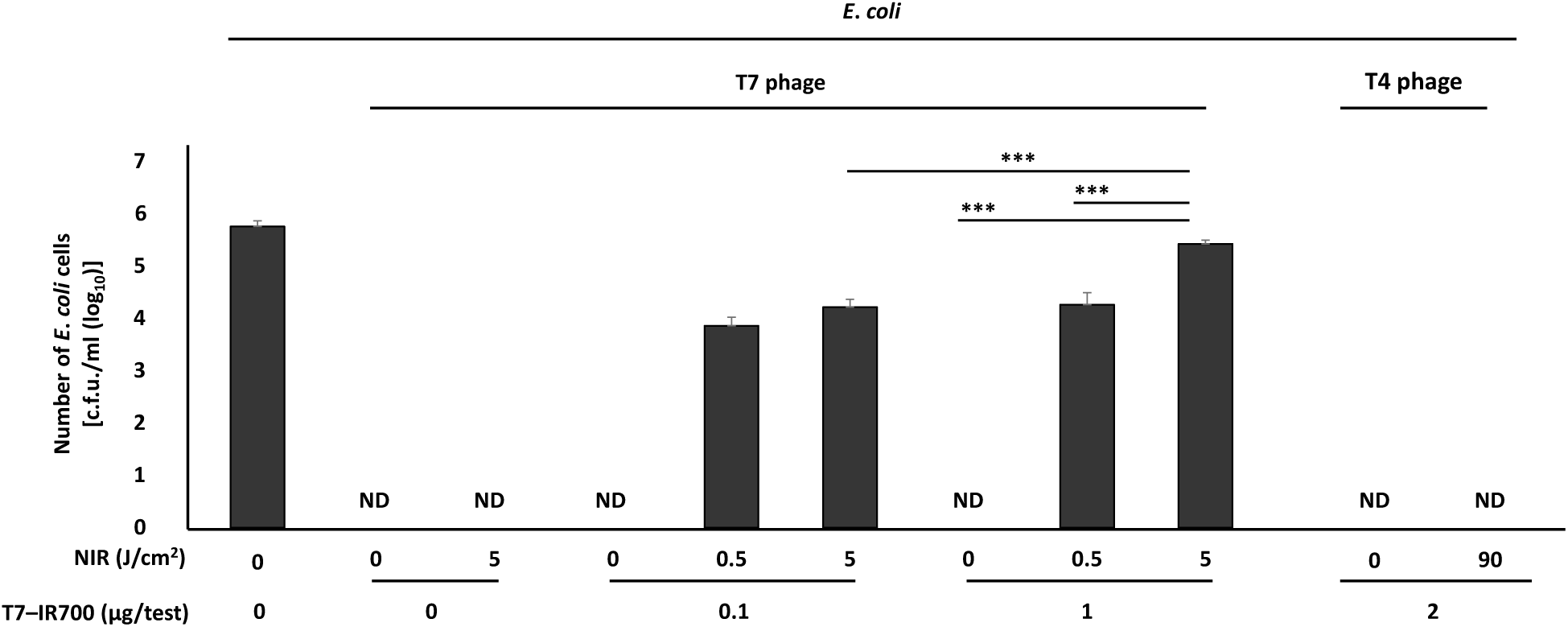
Antiviral effect of PIAS. T7 and T4 phages (10^9^ plaque forming units/test) were treated with PIAS. Subsequently, the PIAS-treated phage samples were co-cultured with *E. coli* cells, and were cultured on agar plates. *E. coli* colonies were enumerated to evaluate the inactivation of the phage by PIAS. c.f.u., colony forming units; ND, not detected. Data are merged from two different experiments (T7 and T4). Mean values are shown, and error bars represent standard deviation from triplicate samples. Two-way analysis of variant with Sidak test. \*\*\**P* < 0.001.

Taken together, PIAS specifically and promptly eliminated the antibody-targets irrespective of the target’s microbial species or drug-resistance status.

## Discussion

Recently, a research group has developed G0755 (ref. 28), a new class of antibiotics against gram-negative pathogens, which are causative agents of hard-to-treat drug-resistant infections. This is the first time in 50 years a new class of antibiotics has been developed. However, the development of new antibiotics has almost reached a theoretical saturation point in which no new ideas are being produced. Conversely, molecular-targeted drugs and antibodies are being explored as front-line therapy because their specificity can reduce unwanted side-effects. There are several studies using antibodies to target *S. aureus* virulence factors (ref. 29) secreted by a subset of *S. aureus* strains or antibiotic conjugated (rifalog, a RFP derivative)-antibodies to target intracellular pathogen cells (ref. 30).

PIAS, a target-specific anti-infective therapy that can be applied to different microbial pathogens, promptly exhibited a target-selective antimicrobial effect and did not lead to the induction of resistant strains. These properties of PIAS appear to be ideal for anti-infective therapy. However, PIAS has limitations similar to conventional anti-infection therapy using antimicrobials. In closed focus, topical administration of conjugates is needed due to difficulty in their accessibility. Using NIR laser fibres, light can be delivered deeper into tissues or organs through intravascular or endoscopic approaches. However, suitable ways should be chosen depending on the situation. PIAS treatment requires antibodies against specific targets; therefore, if the antibody is available, PIAS could be promptly used to treat infections and this would apply to emerging infectious diseases as well. Bacterial and fungal pathogens recalcitrant to antimicrobials and other microbial pathogens (viruses, protozoa, and parasites) may also be attractive future targets for PIAS.

In conclusion, this study raises the possibility of a new type of anti-infection strategy, contributing to advances in anti-infection strategies.

## Acknowledgements

We thank Prof. Longzhu Cui, Drs. Shinya Watanabe, Kotaro Kiga, Yoshifumi Aiba and staff (Jichi Medical Univ.) for providing bacterial strains and their support. We also thank Yukari Dan and Hidehiro Yamada (Hitachi High-Technologies Co., Japan), Hideki Saito and Prof. Toshiaki Tachibana (Jikei Univ.) for preparing and acquiring SEM images, Naoko Toda and colleagues in our laboratory for technical support, Dr. Wataru Suda (RIKEN) for statistical support in a metagenome analysis, and Profs and Drs. Hirotaka Kanuka, Yuki Kinjo, Yoshinobu Manome, Hisao Tajiri, Masayuki Saruta, Akiko Tajima (Jikei Univ.) and Ichiro Imanishi (TUAT) for their support and comments. This study was partly supported by Grant-in-Aids for Challenging Exploratory Research (JSPS KAKENHI 26670487 to M.M.) and Young Scientists (B) (JSPS KAKENHI 17K16233 to K.I.), Takeda Science Foundation (M.M.), and The Jikei University Research Fund (M.M. and T.I.).

## Author contributions

M.M. designed the study and performed *in vitro* and *in vivo* experiments. K.I. performed *in vitro* experiments. T.N. performed *in vivo* experiments. H.M. provided experimental animals. Y.M. and H.K. supported the study and provided comments. T.I. designed the study, performed *in vitro* experiments and wrote the manuscript.

## Supplementary information

### Materials and Methods

#### Microbes and culture conditions

Various strains of *Staphylococcus aureus* (Supplementary Fig. 6), including MSSA (JKmsSA1 and JCM2874), MRSA (JKmrSA1, N315 and USA300), mupirocin-resistant MRSA (JKmmrSA1 resistant to mupirocin 62.5 µg/ml), and VISA (MU3 and MU50), were used. *Staphylococcus epidermidis* JCM2414, *Staphylococcus microti* (an isolate from the nares of a cotton rat), *Corynebacterium tuberculostearicum* JKCt1 (an isolate from the nares of a healthy volunteer), *Candida albicans* [JCM1452, JCM2085 with resistance to azole drugs (fluconazole, itraconazole, and voriconazole), and JKfarCA1 with resistance to flucytosine 32 µg/ml], *Candida stellata* NBRC0703, *Saccharomyces cerevisiae* DYSc, and *Escherichia coli* DH5α were also used. Further, T7 (NBRC20007) and T4 (NBRC20004) phages were employed. Trypticase soy broth (TSB), brain– heart infusion broth (BHI), LB medium (LB), mannitol salt agar with egg yolk (MSA), TSA, and OPAII *Staphylococcus* agar were obtained from Becton, Dickinson and Company. TSB with 0.5% glucose, Sabouraud broth, and equine defibrinated blood (Kohjin Bio Co., Ltd., Saitama, Japan) were also used. CROMagar was obtained from Kanto Chemical Co., Inc. (Tokyo, Japan).

In addition to cells in the exponential phase, microbial cells in the stationary phase (including persister cells that were recalcitrant to antibacterial agents) (ref. 11) were used. Microbial cells were cultured at 37 °C with shaking for 16 h (stationary phase). To obtain microbial cells at the exponential phase, cells were harvested at approximately OD 0.4, determined with a photometer (Mini Photo 518R; TAITEC Co., Saitama, Tokyo).

#### Reagents

Anti-*S. aureus* monoclonal antibody (mAb) (SA; clone Staph12-569.3, murine IgG3), which recognizes the peptidoglycan of *S. aureus*, was purchased from QED Bioscience Inc. (San Diego, CA, USA). Anti-*C. albicans* mAb (CA; clone MC3, murine IgG3), which recognizes the putative β-1, 2-mannan epitope in cell wall mannoproteins and phospholipomannans of *C. albicans*, was purchased from ISCA Diagnostics Ltd. (Exeter, UK). Anti-T7 phage mAb (T7•Tag Antibody, murine IgG2b) directed against the 11 amino-acid gene 10 leader peptide (MetAlaSerMetThrGlyGlyGlnGlnMetGly) of T7 phage was purchased from Merck KGaA (Darmstadt, Germany). Anti-human epidermal growth factor receptor 2 mAb, trastuzumab (Herceptin, humanized IgG1) was purchased from Chugai Pharmaceutical Co. Ltd. (Tokyo, Japan). IRDye700DX (IR700) was purchased from LI-COR Biosciences (Lincoln, NE, USA). RPMI 1640 Medium without phenol red was purchased from Thermo Fisher Scientific (Tokyo, Japan).

#### Synthesis and purification of IR700-conjugated mAb

IR700-conjugating mAb was synthesized as previous described (refs. 6). Briefly, mAb (1.0 mg, 6.8 nmol) was incubated with IR700 (66.8 µg, 34.2 nmol) in 0.1 M Na_2_HPO_4_ (pH 8.5) at room temperature for 1 h. The mixture was purified with a Sephadex G50 column (PD-10; GE Healthcare, Piscataway, NJ, USA). To confirm the number of fluorophore molecules conjugated to each mAb molecule, the concentrations of protein and IR700 were measured spectroscopically based on their absorption at 280 nm and 689 nm, respectively (UV-1800; Shimadzu Corp., Kyoto, Japan). In the present study, conjugates that had approximately three IR700 molecules per mAb molecule were used.

#### Binding of mAb–IR700 conjugates to microbial cells

mAb–IR700 conjugate (1 µg) was added to approximately 1 × 10^5^ colony-forming units (c.f.u.) of bacterial suspension (total volume of 100 µL) and incubated 1 h at 4 °C followed by cell washing with RPMI. The fluorescence of IR700 was measured with a flow cytometry analyzer (MACSQant analyzer; Miltenyi Biotec, Bergisch Gladbach, Germany) and fluorescence microscopy (IX73; Olympus, Tokyo, Japan) with the following filter settings: 608–648-nm excitation filter and 672–712-nm emission filter. To confirm the target specificity of mAb–IR700 conjugate, unconjugated mAb was added before mAb–IR700 treatments.

#### Scanning electron microscopy (SEM)

SEM analyses were performed to detect mAb binding to bacterial cells. The SA–IR700 conjugate-treated *S. aureus* JKmrSA1 cell suspension was washed with phosphate-buffered saline (PBS) containing 1% bovine serum albumin (BSA) and re-suspended in PBS containing 1% BSA. Colloidal gold (12 nm) goat anti-mouse IgG (Jackson Immunoresearch Laboratories, Inc. West Grove, PA, USA) (1 : 25) was added to the solution and incubated for 1 h at room temperature (25 °C). The mixture was dropped onto a nano-percolator to remove unbound antibody and washed with PBS. The samples were analyzed using SEM (SU8010; Hitachi, Ltd., Tokyo, Japan). PIAS-treated cells were also subjected to SEM analysis.

#### mAb–IR700 conjugates mediate PIAS *in vitro*

mAb–IR700 conjugates (0.01 to 20 µg) were added to approximately 1 × 10^5^ c.f.u. of microbial suspension (100 µL of total volume) and incubated for 1 h at 4 °C in the dark. Microbial cells were then irradiated with near-infrared (NIR) illumination (5 to 90 J/cm^2^) using a light-emitting diode releasing light at 670–710 nm (L690-66-60; Epitex Inc., Kyoto, Japan) (refs. 5, 31, 32). A power density of 24 mW/cm^2^ was measured with an optical power meter (PM 100, Thorlabs, Newton, NJ, USA). Serially diluted samples were plated on agar plates for overnight culture to determine microbial viability.

#### PIAS for bacterial viruses

T7 and T4 phages (10^9^ plaque-forming unit/test) were treated with T7–IR700 conjugate (0.1–2 µg/test) at 4 °C for 1 h and then divided into two samples. One sample was treated with NIR illumination while the other was left untreated. Both samples were co-cultured with *E. coli* DH5α cells at 37 °C for 20 min. After co-culture, the samples were cultured on agar plates, and bacterial colonies were enumerated.

#### *In vivo* studies

Animal studies were performed in accordance with the guidelines established by the Animal Care Committee at the Jikei University School of Medicine. All *in vivo* experiments were performed under anesthesia using isoflurane.

#### PIAS in an *in vivo* colonization model

The cotton rat nasal colonization model (ref. 16) was used to determine the feasibility of PIAS *in vivo*. Six-to ten-week-old cotton rats (*Sigmodon hispidus*) were obtained from the Animal Research Center, University of Occupational and Environmental Health School of Medicine (Fukuoka, Japan). All rats were allowed to acclimatize and recover from shipping-related stress for one week and were kept under a controlled light/dark cycle (12 : 12 h) before the experiments were performed. Ten microliters of a suspension of 1 × 10^6^ c.f.u. of *S. aureus* JKmrSA1 cells was instilled in both nares of cotton rats. The animals were then kept supine for approximately 1 h for recovery from anesthesia. Seven days after intranasal instillation, animals were randomized into five groups, with at least three rats per group, which were treated as follows: (i) intranasal administration of PBS (10 µL); (ii) NIR laser illumination (50 J/cm^2^); (iii) intranasal administration of PBS (10 µL), followed by NIR laser illumination (50 J/cm^2^); (iv) intranasal administration of SA–IR700 conjugates (5 µg, 10 µL), followed by NIR laser illumination (10 J/cm^2^); or (v) intranasal administration of SA–IR700 conjugates (5 µg, 10 µL), followed by NIR laser illumination (50 J/cm^2^).

All procedures were performed under anesthesia and NIR laser illumination was performed 1 h after intranasal administration of SA–IR700 or PBS. NIR laser illumination was performed with a 690-nm continuous-wave laser at a power density of 330 mW/cm^2^ (ML6540-690; Modulight, Inc., Tampere, Finland). After NIR laser treatment, the animals were killed and the face, specifically the exterior of the nose, was carefully disinfected with 70% alcohol. The anterior nares were then harvested by dissecting the nose. Harvested samples were collected in PBS with Tween 20 and vortexed at maximum speed for 1 min. Serially diluted samples were plated on agar plates for overnight culture to determine bacterial viability. The absence of *S. aureus* contamination was confirmed for all animals before their use in these experiments.

#### PIAS in an intraperitoneal infection model

A mouse model of intraperitoneal infection (ref. 17) was used. Five-to seven-week-old female specific-pathogen-free BALB/c mice were obtained from Clea Japan (Tokyo, Japan). Mice were intraperitoneally injected with 100 µL suspension of 2 × 10^8^ c.f.u. of *S. aureus* JKmrSA1 cells and treated with PIAS or antibiotics (VCM or RFP and VCM). (i) PIAS-treated group (*n* = 8): SA–IR700 (50 µg/mouse) and NIR illumination (50 J/cm^2^) from outside the body encompassing the entire abdomen. (ii) PIAS-control group (*n* = 8): SA–IR700 (50 µg/mouse) without NIR illumination. (iii) VCM-treated group (*n* = 3): VCM in 10 mg/kg. (iv) RFP and VCM-treated group (*n* = 8): RFP in 100 mg/kg and VCM in 50 mg/kg. (v) Control (*n* = 3): 100 µL of PBS. SA–IR700 and VCM or RFP were intraperitoneally or perorally administered, respectively.

Treatments were performed within 1 h after infection once during the experimental period while only VCM was administered twice. Survival and adverse events were monitored via a once-daily assessment for seven days. Faeces of PIAS and antibiotic (VCM and RFP)-treated mice and those of non-treated mice were used for 16S-targeting metagenome analysis (Fig. 2E).

#### PIAS in a murine thigh infection model

A mouse model of thigh infection (ref. 18) was used. A 100 µL suspension of 1 × 10^7^ c.f.u. of *S. aureus* JKmrSA1 cells were injected into the right thigh muscle of six-to seven-week old female mice and treated with or without PIAS (*n* = 8 each group). PIAS conditions: SA–IR700 (50 µg/mouse) and NIR illumination (50 J/cm^2^) from outside the body encompassing the right thigh. Control: SA–IR700 (50 µg) without NIR illumination. One day after treatment, mice were killed, and the right thighs were dissected out. Thigh samples were collected in PBS with Tween 20 and homogenized manually. The homogenized samples were cultured on OPAII to evaluate the bactericidal effect of PIAS on the pathogen cells in the thigh. Moreover, thigh samples 7 days after treatment were employed to assess the pathology. Macroscopic findings were confirmed histologically as appropriate with hematoxylin-eosin staining.

#### Metagenome analysis (refs. 19, 20)

DNA was extracted from faecal pellets and PCR was performed using 27Fmod 5′-AGRGTTTGATYMTGGCTCAG-3′ and 338R 5′-TGCTGCCTCCCGTAGGAGT-3′ to the V1–V2 region of the 16S rRNA gene. The 16S metagenomic sequencing was performed using MiSeq according to the Illumina protocol. Two paired-end reads were merged using the fastq-join program based on overlapping sequences. Reads with an average quality value of < 25 and inexact matches to both universal primers were filtered out. Filter-passed reads were used for further analysis after trimming off both primer sequences. For each sample, 3,000 quality filter-passed reads were rearranged in descending order according to the quality value and then clustered into OTUs with a 97% pairwise-identity cutoff using the UCLUST program (Edgar 2010) version 5.2.32 (https://www.drive5.com). Taxonomic assignment of each OTU was made via searching by similarity against the RDP and the NCBI genome database using the GLSEARCH program.

#### Statistical analyses

Mean ± standard deviation values were calculated from a minimum of three samples. Calculations and statistical analyses were performed using Prism software (ver. 8, GraphPad Software Inc., La Jolla, CA, USA) and Excel software (Microsoft, Redmond, WA, USA). The two-sided Student’s *t*-test was used to determine the differences in bacterial count between the two treatment groups. One-way analysis of variance (ANOVA) with Dunnett’s test or with Sidak test, or two-way ANOVA with Sidak test was performed for multiple group comparisons. Kaplan Meier survival curve was assessed using a log-rank (Mantel-Cox) test. In a metagenome analysis, ANOVA was performed for multiple group comparisons, and any significant differences were evaluated using two-stage step-up method of Benjamini, Krieger and Yekutieli (ref. 33). Results were considered to be statistically significant at a p value < 0.05.

**Supplementary Figure 1.**
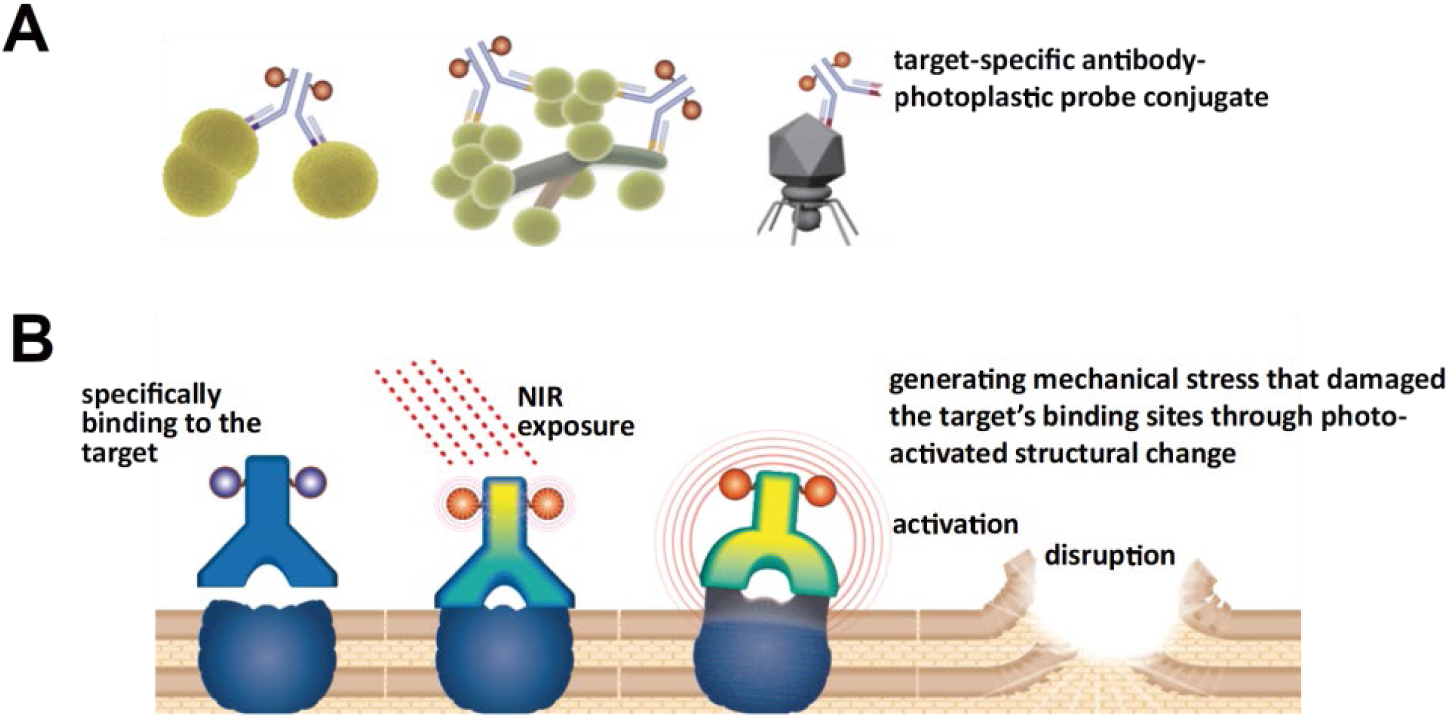
Model of binding of mAb-photoplastic probe conjugates to the targets and the proposed mechanism of PIAS. **A**. Anti-pathogen antibody-photoplastic probe IR700 conjugates specifically bind to the antibody targets. **B**. Scheme indicates the proposed mechanism of PIAS. The recent study (ref. 4) reported that NIR induces an axial ligand-releasing reaction, which dramatically alters hydrophilicity of IR700; NIR illumination causes physical changes in the shape of antibody antigen complexes once the conjugate is bound to its target, inducing mechanical stress to the surface epitopes and disrupting it. Our study indicated that PIAS specifically and promptly damages target cells when conjugate binding is followed by NIR illumination and suggested that PIAS can act on pathogens differently than antimicrobials (ref. 14) or photodynamic therapy (ref. 15).

**Supplementary Figure 2.**
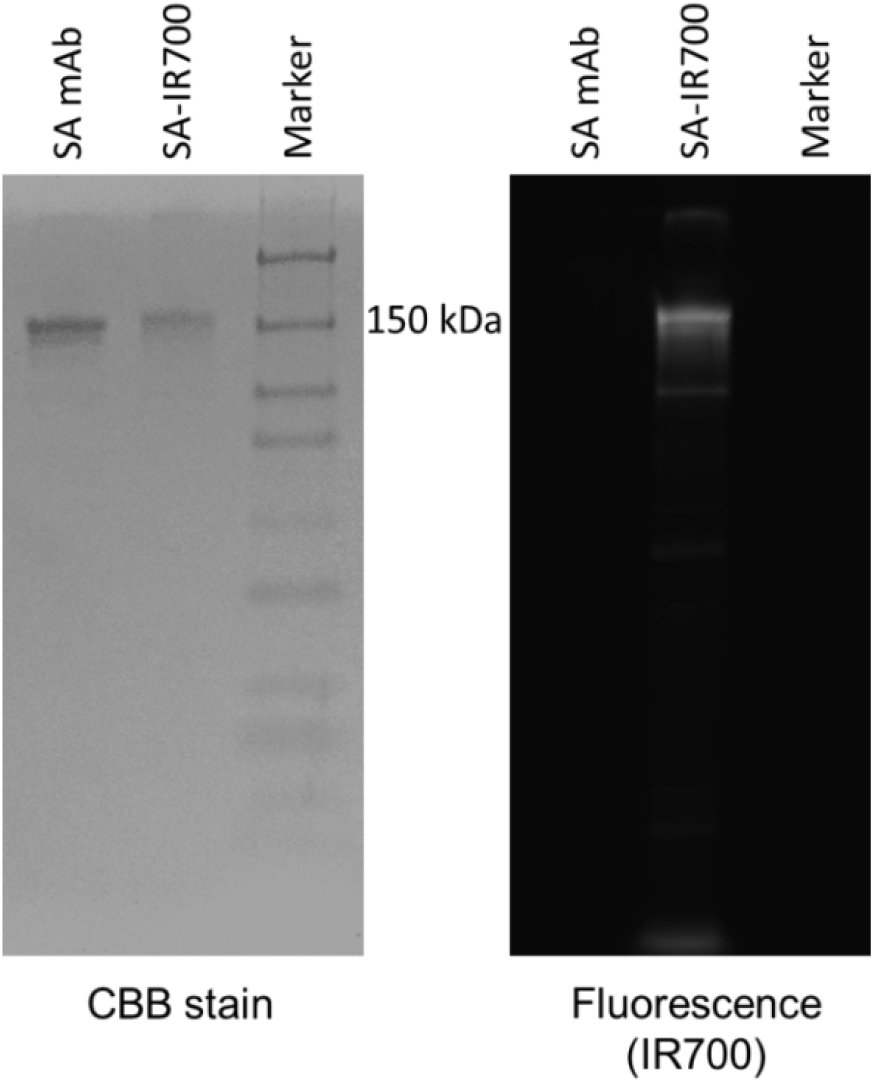
IR700 fluorescence derived from mAb–IR700 conjugates. SA–IR700 conjugates and SA mAb were subjected to non-reducing SDS-PAGE. Subsequently, the gel was stained using Coomassie Brilliant Blue (CBB) to obtain IR700 fluorescence. IR700 fluorescence was observed corresponding with the band of SA–IR700 conjugates in the gel.

**Supplementary Figure 3.**
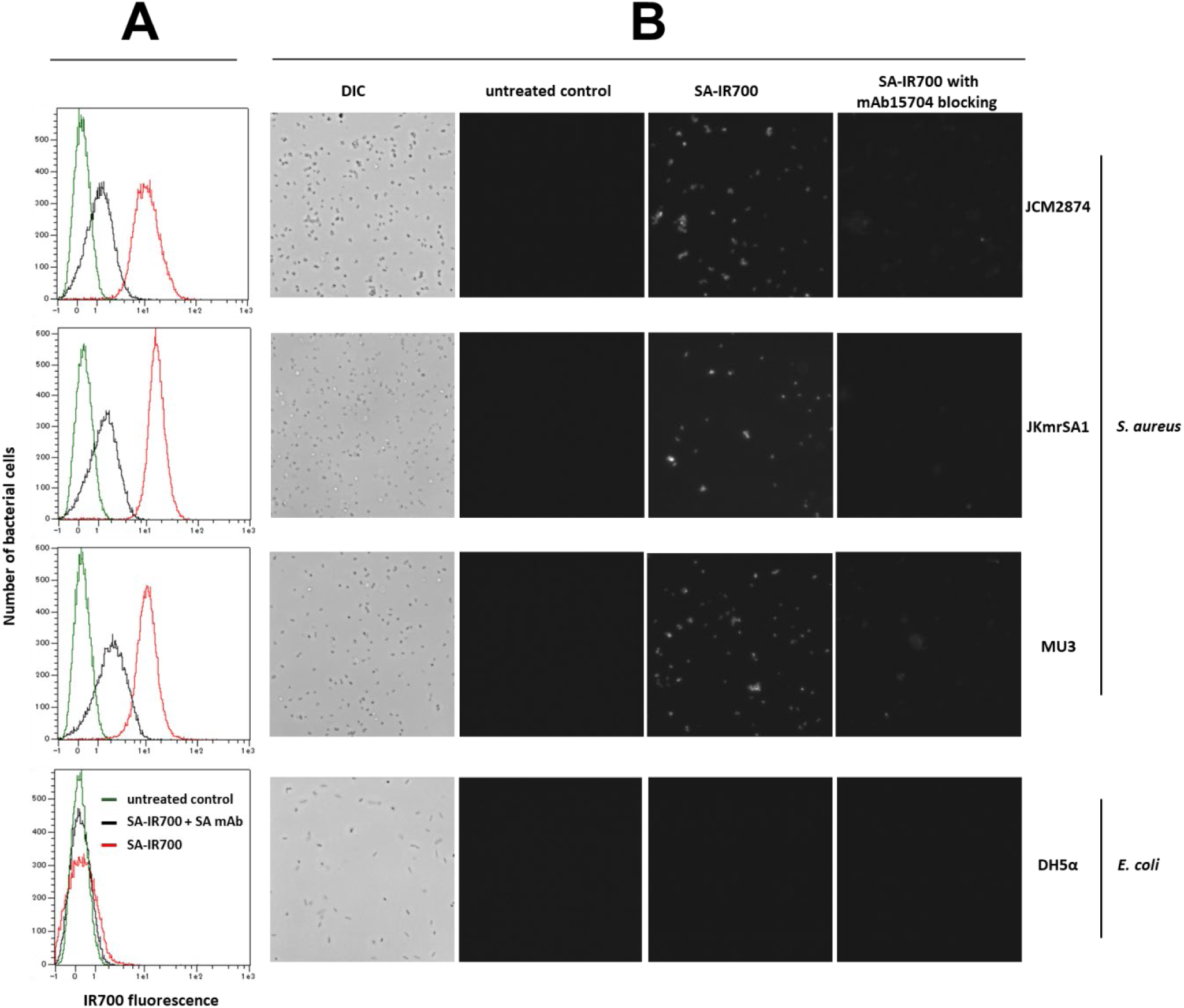
Binding of mAb-conjugates to the target cells. **A** and **B**. Binding ability of SA–IR700 conjugates toward *S. aureus* and *E. coli* cells was examined using flow cytometry (**A**) and fluorescence microscopy (**B**). SA–IR700 conjugates (2 μg/test) and SA mAb (20 μg/test) were used. **A.** Bacterial cells of various *S. aureus* strains treated with SA–IR700 conjugates or SA–IR700 conjugates with SA mAb blocking were subjected to flow cytometric analysis. The Y-axis indicates the number of bacterial cells, and the X-axis indicates fluorescence of IR700. **B.** Bacterial cells treated with SA–IR700 conjugates or SA–IR700 conjugates with SA mAb blocking were subjected to fluorescence or differential interference contrast (DIC) microscopic analyses. Bright spots indicate fluorescence derived from SA–IR700 conjugates. Representative images are shown.

**Supplementary Figure 4.**
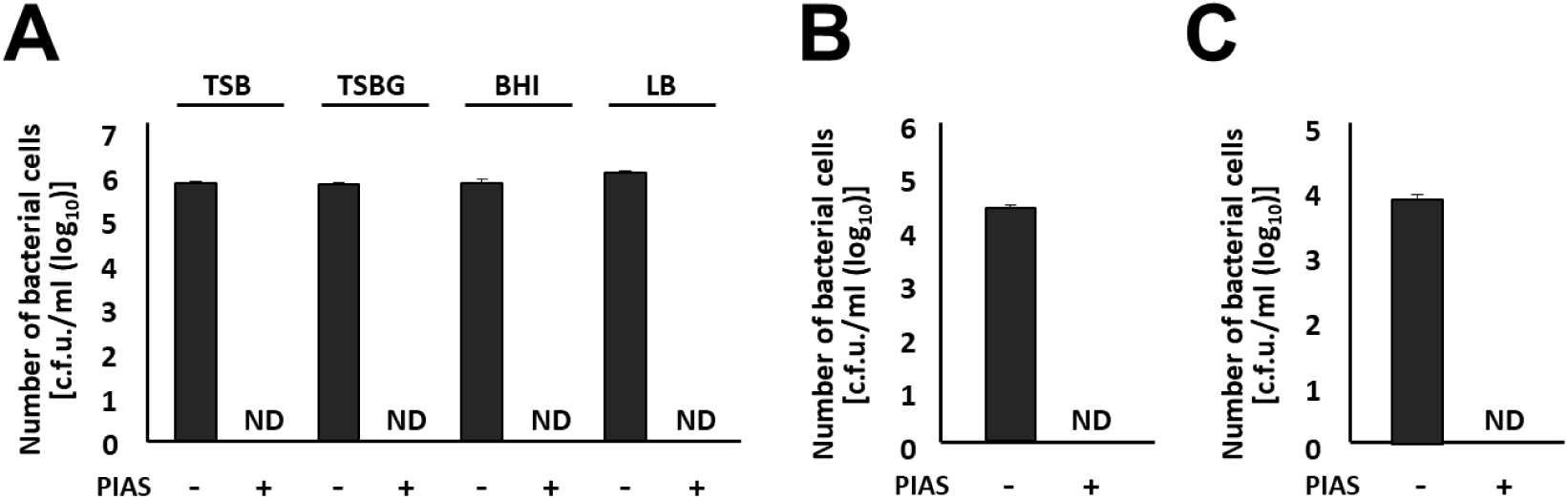
Effect of PIAS on *S. aureus* under various conditions. Bacterial cells of *S. aureus* JKmsSA1 were subjected to the following experiments. **A.** Bacterial cells cultured in various media were subjected to PIAS. PIAS conditions: SA–IR700, 1 µg/test; NIR illumination, 15 J/cm^2^. **B.** Bacterial cells were suspended in equine defibrinated blood and then subjected to PIAS. PIAS conditions: SA–IR700, 4 µg/test; NIR illumination, 30 J/cm^2^. **C.** Bacterial cells (1 × 10^7^ cells/test) treated with SA–IR700 conjugates (30 µg/test) were added in 3T3 cells (confluent cells in 35 mm dish) and co-cultured at 37 °C under 5% CO_2_ for 30 min (refs. 30, 34). After co-culture, cells were rinsed thrice with 1 mL of RPMI, and then extracellular *S. aureus* was lysed with 1 mL of lysostaphin (FUJIFILM Wako Pure Chemical Co., Tokyo, Japan) solution (1 mg/ml) for 30 min (ref. 34). Cells were rinsed five times with 1 mL of RPMI and treated by illumination to NIR. After treatment, the bacterial samples were homogenized and cultured on agar plates. The bactericidal effect of PIAS was evaluated using the colony-counting method. c.f.u., colony-forming units; ND, not detected. Mean values are shown (*n* = 3). Error bars represent standard deviation.

**Supplementary Figure 5.**
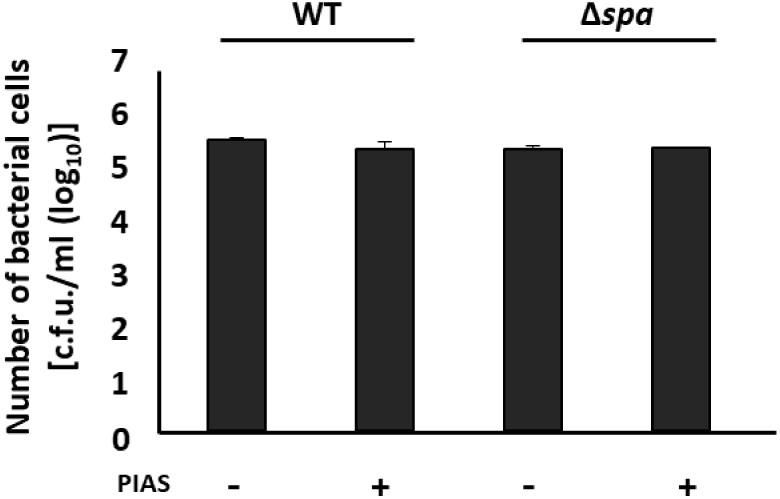
Contribution of protein A on the bactericidal effect of PIAS for *S. aureus* cells. Bacterial cells of Δ*spa* cells and wild type (RN4220) were subjected to PIAS, and then the bacterial samples were cultured on agar plates. The bacterial viability was evaluated using the colony counting method. PIAS conditions: SA–IR700, 2 µg/test; NIR illumination, 90 J/cm^2^. c.f.u., colony forming units. Mean values are shown (*n* = 3). Error bars represent standard deviation.

**Supplementary Figure 6.**
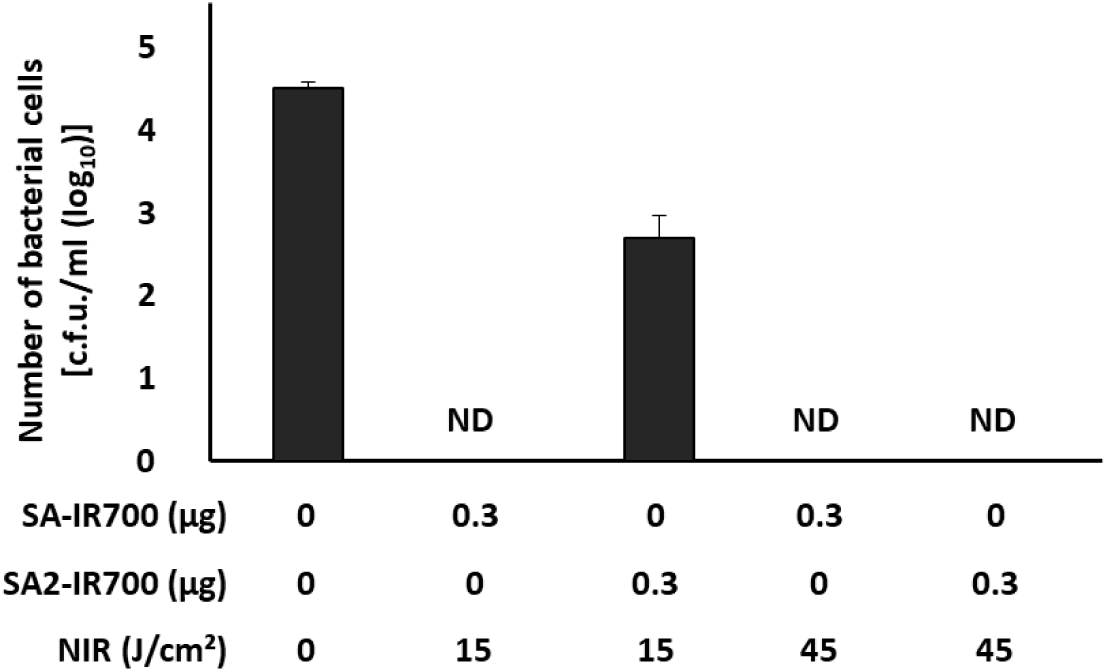
Bactericidal effect of PIAS with an anti-*S. aureus* antibody against different epitopes. Bacterial cells of *S. aureus* JKmsSA1 were subjected to PIAS using the SA mAb [from clone Staph12-569.3 (SA) or from clone Staph11-232.3 (SA2)]–IR700 conjugate; SA2 was only used in this experiment. After PIAS, the bacterial cells were cultured on agar plates, and colonies were enumerated to evaluate the bactericidal effect. c.f.u., colony-forming units; ND, not detected. Mean values are shown (*n* = 3). Error bars represent standard deviation.

**Supplementary Figure 7.**
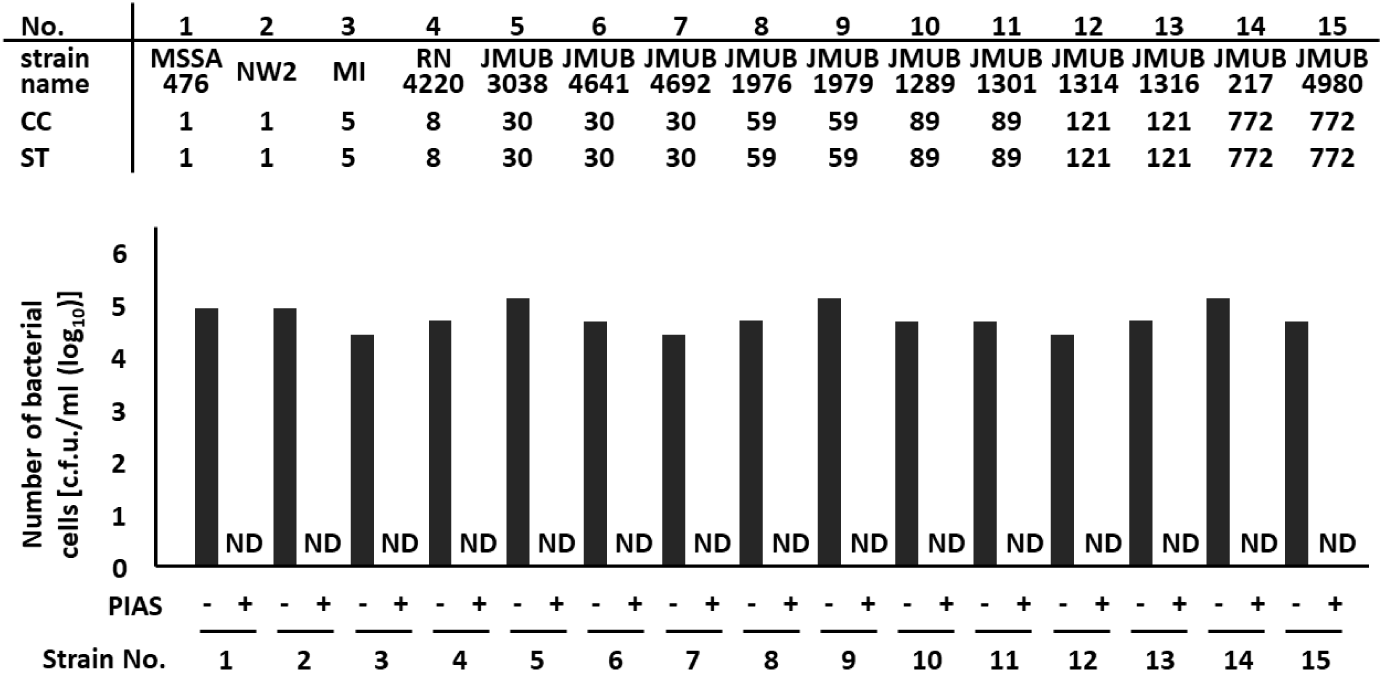
Bactericidal effect of PIAS on various *S. aureus* strains. A diverse panel of *S. aureus* sequence types was tested. Bacterial cells (approximately 10^5^ c.f.u./test) were subjected to PIAS and cultured on agar plates to evaluate the bactericidal effect of PIAS on various *S. aureus* strains. PIAS conditions: SA–IR700, 2 µg/test; NIR illumination, 90 J/cm^2^. c.f.u., colony-forming units; ND, not detected.

**Supplementary Figure 8.**
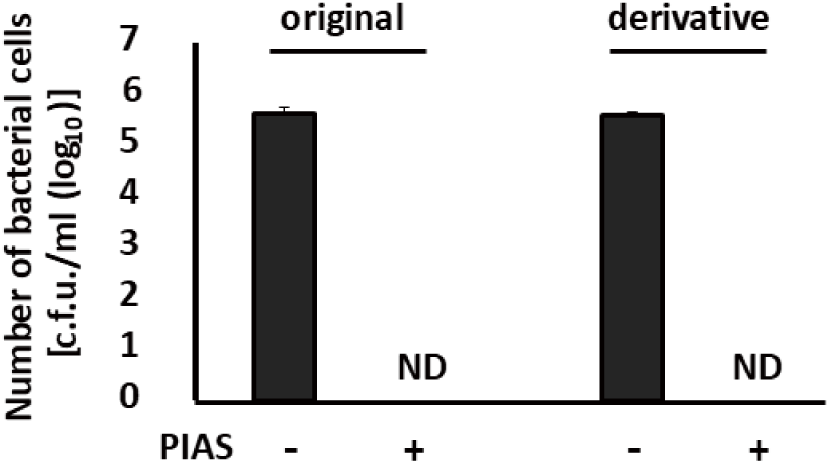
Follow-up experiment for the acquisition of PIAS-resistance. Bacterial cells of *S. aureus* JKmsSA1 (original) and its derivative strain after 30 passages (derivative) were subjected to PIAS. After PIAS, the bacterial cells were cultured on agar plates, and colonies were enumerated to evaluate the bactericidal effect. PIAS conditions: SA–IR700, 2 µg/test; NIR illumination, 90 J/cm^2^. c.f.u., colony-forming units; ND, not detected. Mean values are shown (*n* = 3). Error bars represent standard deviation.

**Supplementary Figure 9.**
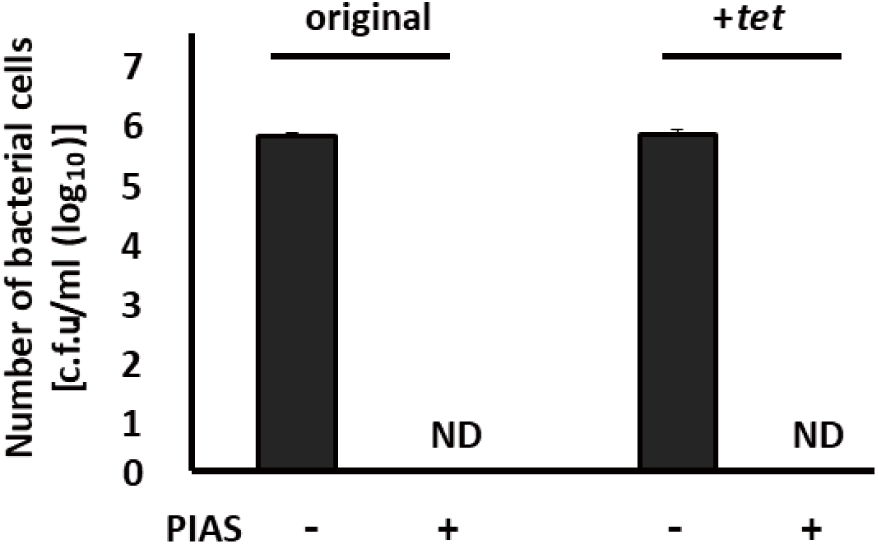
Effect of PIAS on *S. aureus* harboring a plasmid encoding the tetracycline-resistance gene. Bacterial cells of *S. aureus* JKmsSA1 (original) and its derivative strain JKmsSA1 harboring a plasmid encoding the tetracycline-resistance gene (+ *tet*) were subjected to PIAS. PIAS conditions: SA–IR700, 2 µg/test; NIR illumination, 90 J/cm^2^. c.f.u., colony-forming units; ND, not detected. Mean values are shown (*n* = 3). Error bars represent standard deviation.

**Supplementary Figure 10.**
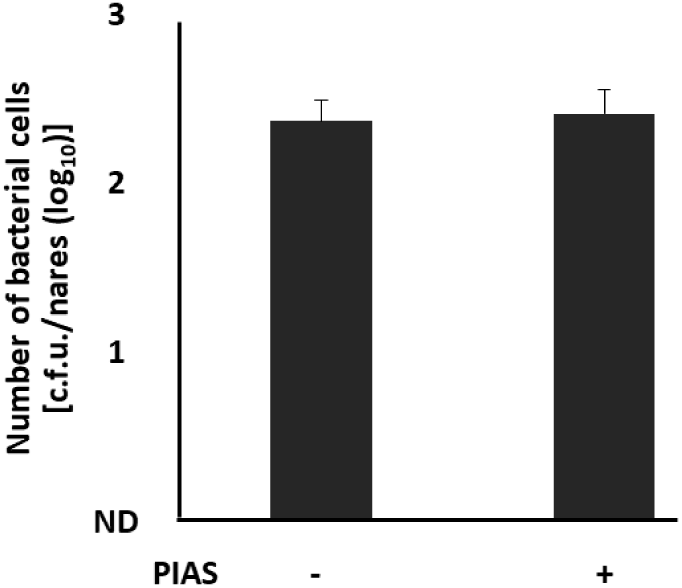
Effect of PIAS on the rat normal flora, including *S. microti*. Animals colonized by MRSA JKmrSA1 were subjected to PIAS using SA-IR700 conjugate (2 µg/test). Subsequently, the samples were cultured on MSA. c.f.u., colony forming units. Mean values are shown (*n* = 3). Error bars represent standard deviation.

**Supplementary Figure 11.**
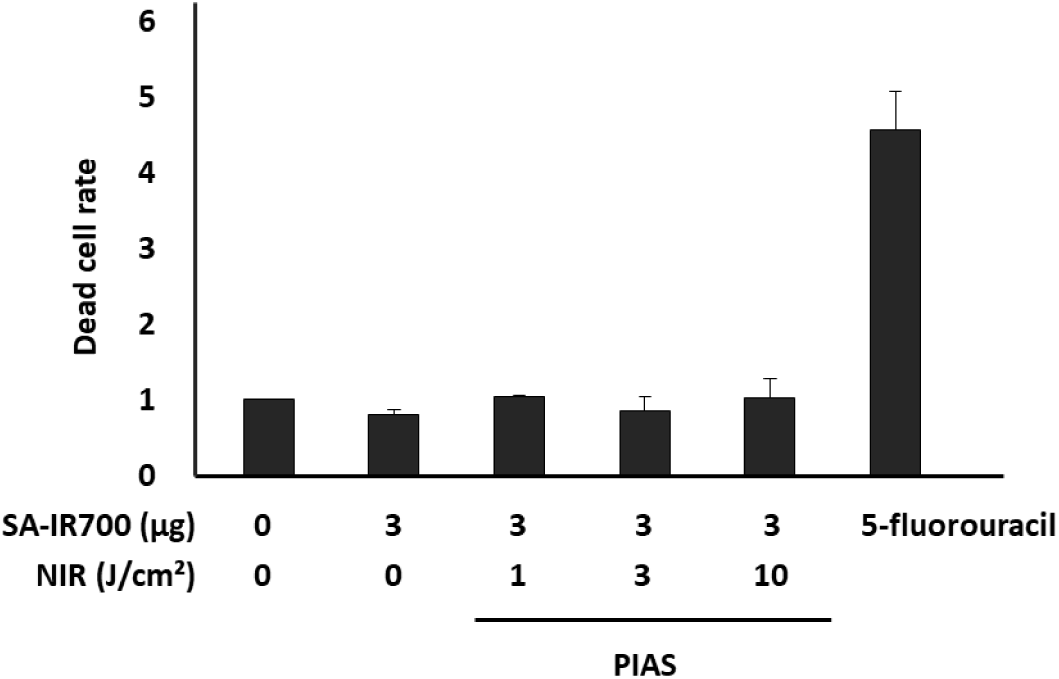
Effect of PIAS on the non-targeted fibroblast cells. Cytotoxic effect of PIAS using SA–IR700 conjugates on the non-targeted fibroblast cells was investigated. Cells of the fibroblast cell line 3T3 were treated with SA– IR700 for 3 h and then exposed to NIR or with 5-fluorouracil (5 µg/mL) for 24 h. The cytotoxicity was evaluated using a live/dead assay. Columns indicate mean values normalized by control (SA-IR700; 0 µg; NIR 0 J/cm_2_). Error bars represent standard error from triplicate samples.

**Supplementary Figure 12.**
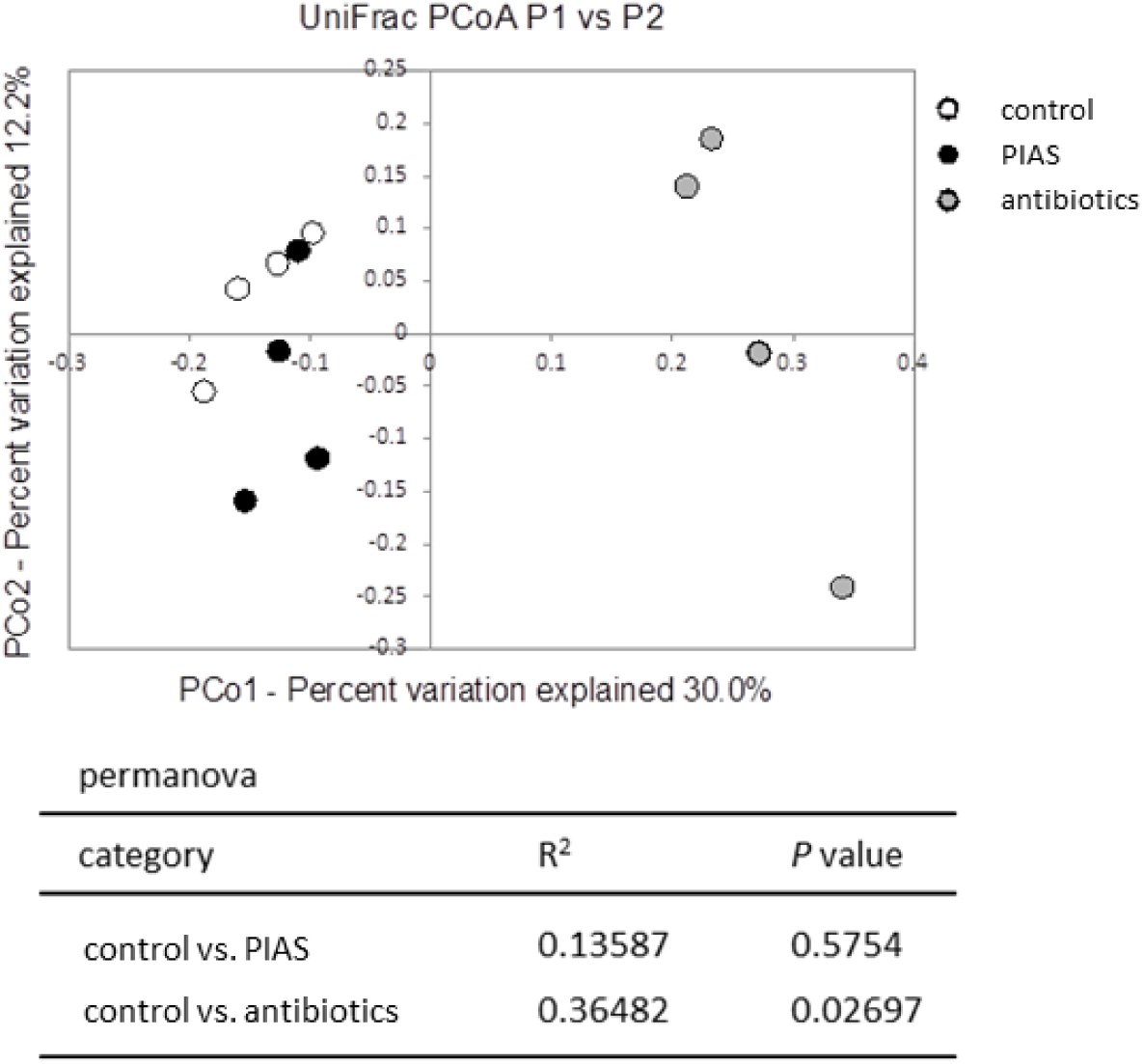
Principal coordinates analysis (PCoA) of UniFrac distances of 16S rRNA genes. PCoA was performed on unweighted UniFrac distance to compare test samples using gene sequence data (ref. 35). control, PBS; antibiotics, VCM and REF.

## References

1. Laxminarayan R, Matsoso P, Pant S, Brower C, Røttingen JA, Klugman K, Davies S. Access to effective antimicrobials: a worldwide challenge. Lancet 2016;387:168–175.

2. Levy SB, Marshall B. Antibacterial resistance worldwide: causes, challenges and responses. Nat. Med. 2004;10:S122–129.

3. Aminov RI. A brief history of the antibiotic era: lessons learned and challenges for the future. Front. Microbiol. 2010;1:134.

4. Lewis K. Antibiotics: Recover the lost art of drug discovery. Nature 2012;485:439–440.

5. Sato K, et al. Photoinduced ligand release from a silicon phthalocyanine dye conjugated with monoclonal antibodies: A mechanism of cancer cell cytotoxicity after near-infrared photoimmunotherapy. ACS Cent. Sci. 2018;4:1559–1569.

6. Mitsunaga M, et al. Cancer cell-selective *in vivo* near infrared photoimmunotherapy targeting specific membrane molecules. Nat. Med. 2011;17:1685–1691.

7. ClinicalTrials.gov Identifier: NCT02422979 (phase I/II) and NCT03769506 (phase III).

8. Lowy FD. *Staphylococcus aureus* infections. N. Engl. J. Med. 12 1998;339:520–532.

9. Chambers HF, Deleo FR. Waves of resistance: *Staphylococcus aureus* in the antibiotic era. Nat. Rev. Microbiol. 2009;7:629–641.

10. Brown AF, Leech JM, Rogers TR, McLoughlin RM. *Staphylococcus aureus* colonization: modulation of host immune response and impact on human vaccine design. Front. Immunol. 2014;4:507.

11. Harms A, Maisonneuve E, Gerdes K. Mechanisms of bacterial persistence during stress and antibiotic exposure. Science 2016;354:aaf4268.

12. Hiramatsu K, et al. Dissemination in Japanese hospitals of strains of *Staphylococcus aureus* heterogeneously resistant to vancomycin. Lancet 1997;350:1670–1673.

13. Grov A, Myklestad B, Oeding P. Immunochemical studies on antigen preparations from *Staphylococcus aureus*. Isolation and chemical characterization of antigen A. Acta Pathol. Microbiol. Scand. 1964;61:588–596.

14. McDermott PF, Walker RD, White DG. Antimicrobials: modes of action and mechanisms of resistance. Int. J. Toxicol. 2003;22:135–143.

15. Wainwright M, Maisch T, Nonell S, Plaetzer K, Almeida A, Tegos GP, Hamblin MR. Photoantimicrobials-are we afraid of the light? Lancet Infect. Dis. 2017;17:e49–e55.

16. Kokai-Kun JF. The cotton rat as a model for *Staphylococcus aureus* nasal colonization in humans: cotton rat *S*. aureus nasal colonization model. Methods Mol. Biol. 2008;431:241–254.

17. Liu CI, et al. A cholesterol biosynthesis inhibitor blocks *Staphylococcus aureus* virulence. Science 2008;319:1391–1394.

18. Hertlein T, et al. Visualization of abscess formation in a murine thigh infection model of *Staphylococcus aureus* by ^19^F-magnetic resonance imaging (MRI). PLoS One 2011;6:e18246.

19. Tanoue T, et al. A defined commensal consortium elicits CD8 T cells and anti-cancer immunity. Nature 2019;565:600–605.

20. Iwasawa K, et al. Dysbiosis of the salivary microbiota in pediatric-onset primary sclerosing cholangitis and its potential as a biomarker. Sci. Rep. 2018;8:5480.

21. Kim S, Covington A, Pamer EG. The intestinal microbiota: Antibiotics, colonization resistance, and enteric pathogens. Immunol. Rev. 2017;279:90–105.

22. Kalghatgi S, et al. Bactericidal antibiotics induce mitochondrial dysfunction and oxidative damage in mammalian cells. Sci. Transl. Med. 2013;5:192ra85.

23. Parker JC, McCloskey JJ, Knauer KA. Pathobiologic features of human candidiasis. A common deep mycosis of the brain, heart and kidney in the altered host. Am. J. Clin. Pathol. 1976;65:991–1000.

24. Ostrosky-Zeichner L, Casadevall A, Galgiani JN, Odds FC, Rex JH. An insight into the antifungal pipeline: selected new molecules and beyond. Nat. Rev. Drug Discov. 2010;9:719–727.

25. Doermann AH. Lysis and lysis inhibition with *Escherichia coli* bacteriophage. J. Bacteriol. 1948;55:257–276.

26. Kutter E, et al. Phage therapy in clinical practice: treatment of human infections. Curr. Pharm. Biotechnol. 2010;11:69–86.

27. Dedrick RM, et al. Engineered bacteriophages for treatment of a patient with a disseminated drug-resistant *Mycobacterium abscessus*. Nat. Med. 2019;25:730–733.

28. Imai Y, et al. A new antibiotic selectively kills Gram-negative pathogens. Nature 2019;576:459–464.

29. Chan R, et al. Identification of biologic agents to neutralize the bicomponent leukocidins of *Staphylococcus aureus*. Sci. Transl. Med. 2019;11:eaat0882.

30. Lehar SM, et al. Novel antibody–antibiotic conjugate eliminates intracellular *S. aureus*. Nature 2017;527:323–328.

## Supplementary references

31. Mitsunaga M, Nakajima T, Sano K, Choyke PL, Kobayashi H. Nearinfrared theranostic photoimmunotherapy. (PIT): repeated exposure of light enhances the effect of immunoconjugate. Bioconjug. Chem. 201;23:604–609.

32. Ito K, Mitsunaga M, Nishimura T, Kobayashi H, Tajiri H. Combination photoimmunotherapy with monoclonal antibodies recognizing different epitopes of human epidermal growth factor receptor 2: an assessment of phototherapeutic effect based on fluorescence molecular imaging. Oncotarget 2016;7:14143–14152.

33. Benjamini Y, Krieger AM, Yekutieli D. Adaptive linear step-up procedures that control the false discovery rate. Biometrika 2006;93:491–507.

34. Usui A, Murai M, Seki K, Sakurada J, Masuda S. Intracellular localization of *Staphylococcus aureus* within primary cultured mouse kidney cells. Microbiol. Immunol. 1992;36:545–550 (1992).

35. Lozupone C, Lladser ME, Knights D, Stombaugh J, Knight R. UniFrac: an effective distance metric for microbial community comparison. ISME J. 2011;5:169–172.

